# ATP-driven membrane binding and polymerization of bacterial actin MreB promotes local membrane fluidization

**DOI:** 10.64898/2026.02.13.705564

**Authors:** Ingrid E. Adriaans, Ana Álvarez-Mena, Céline Dinet, Estelle Morvan, Kee Siang Lim, Erick J. Dufourc, Richard Wong, Arnaud Chastanet, Alphée Michelot, Rut Carballido-López, Birgit Habenstein

**Author notes:** corresponding author: R. Carballido-López; B. Habenstein. these authors contributed equally (Ingrid E. Adriaans, Ana Álvarez-Mena).

## Abstract

The bacterial actin homologue MreB plays a key role in rod cell shape determination. We recently showed that MreB from the Gram-positive bacterium *Geobacillus stearothermophilus* (MreB^Gs^) polymerizes into straight pairs of protofilaments in the presence of both ATP and a lipid surface. Membrane interaction is thought to be mediated by electrostatic interactions with anionic lipids, with final anchoring relying on two spatially close hydrophobic motifs that protrude from the MreB^Gs^ monomers, forming a putative membrane-insertion domain. Here, we determined the binding properties of ATP and ADP to MreB^Gs^ using fluorescence anisotropy, and monitored ATP-mediated binding and polymer formation on lipid bilayers using liposome binding assays and AFM, respectively. Finally, we used solid-state NMR to visualize the interaction between the membrane and MreB^Gs^ at the atomic level. Our findings reveal that MreB^Gs^ has similar affinity for both ATP and ADP, unlike eukaryotic actin. We also show that monomeric MreB^Gs^ establishes peripheral contacts with the membrane likely through electrostatic interactions, while ATP-induced MreB^Gs^ filaments insert into the lipid bilayer without interfering with the membrane lamellar phase and have a significant local fluidifying effect.

**Statement of significance:** Bacteria rely on the actin-like protein MreB to determine and maintain their cell shape, like actin does in eukaryotic cells. To perform its tightly regulated morphogenetic function, MreB forms membrane-associated nanofilaments *in vivo*, which control the cell wall biosynthetic machinery. We recently demonstrated that, *in vitro*, MreB from the Gram-positive bacterium *Geobacillus stearothermophilus* requires both ATP and lipids to polymerize into pairs of filaments. Here, we show that in the presence of ATP, *Geobacillus* MreB forms membrane-bound filaments that directly impact local membrane fluidity, which could translate into a regulatory effect on cell wall synthetic enzymes. We further reveal that MreB binds both ATP and ADP with similar affinity, unlike eukaryotic actin, which preferentially binds ATP over ADP, and speculate that this could be a mechanism modulating the pool of polymerization-competent MreB in bacteria.

## Introduction

Filament-forming cytoskeletal proteins are a hallmark of cells across all domains of life, orchestrating essential processes such as cell shape determination, mechanical stability, cytokinesis, chromosome segregation, and intracellular transport. In most rod-shaped bacteria, the intracellular actin-like MreB proteins and the extracellular peptidoglycan cell wall are the major determinants of cell shape (1). *In vivo*, MreB forms membrane-associated polymeric assemblies that move around the rod circumference together with proteins of the peptidoglycan elongation machinery, forming the so-called Rod complex (2–4). MreB filaments orient this circumferential motion, thereby directing the insertion of new cell wall material in radial hoops and promoting cylindrical growth (2, 3).

*In vitro*, MreB polymerizes into straight pairs of protofilaments (5). Polymerization of MreB from the Gram-positive *Geobacillus stearothermophilus* (MreB^Gs^), and possibly from other Gram positive bacteria, requires the presence of both lipids and ATP (6). Notably, ATP, but not ADP, promotes both MreB^Gs^ membrane binding and polymerization into the canonical pair of straight protofilaments (6). It was assumed that, like actin molecules, MreB^Gs^ binds both ATP and ADP, and hypothesized that only ATP binding or hydrolysis triggers conformational changes that modulate the affinity of MreB^Gs^ for the membrane (6). ATP-bound MreB^Gs^ membrane binding is mediated by two adjacent hydrophobic regions, the N-terminus and a surface-exposed loop, which together form a putative membrane insertion domain, and is further facilitated by electrostatic interactions between charged residues surrounding these regions and anionic lipids (6). Direct membrane binding, via an amphipathic helix and/or a hydrophobic insertion loop, has also been shown for MreB filaments of several Gram-negative species (6–9). However, the insertion of the MreB molecule into the lipid layer has not yet been demonstrated, and the potential effects of MreB filaments on membrane structure or dynamics remain unexplored. *In vivo* data, using lipid staining and spectrometry approaches, suggested that MreB filaments induce the formation of regions of increased fluidity (RIFs) within the membrane of *Bacillus subtilis* (10). Whether this effect was direct or indirect could not be determined within the complex cellular environment, and the mechanism by which MreB was proposed to create fluid lipid domains remains elusive.

Here, we quantified the binding kinetics of ATP and ADP to MreB^Gs^ using fluorescence anisotropy, and we characterized the binding of ATP-MreB^Gs^ to a lipid bilayer using a combination of liposome-binding assays, Atomic Force Microscopy (AFM) and wideline solid-state NMR (ssNMR). Notably, we found that MreB binds both ATP and ADP with similar affinities -unlike eukaryotic actins- and that ATP promotes the insertion of MreB^Gs^ into the lipid bilayer. This insertion significantly increases the fluidity of the hydrophobic core region of the membrane without interfering with the lamellar membrane phase. Furthermore, inorganic phosphate (Pi) was not detected in our ssNMR measurements, suggesting that membrane-bound MreB filaments are predominantly in either the ATP or ADP-Pi state (6) under these conditions. Together, our findings suggest that modulation of the intracellular ATP/ADP ratio could be a regulatory mechanism for MreB polymerization, and support a model in which ATP-induced MreB^Gs^ filaments locally increase lipid-chain disorder, thereby modulating membrane fluidity at the sites of action of the Rod complex.

## Materials and Methods

### Reagents and materials

1-palmitoyl-d31-2-oleoyl-sn-glycero-3-phosphocoline (^2^H_31_-POPC), 1,2-dioleoyl-sn-glycero-3-phosphocholine (DOPC), and 1,2-dioleoyl-sn-glycero-3-phospho-(1’-rac-glycerol) (DOPG) were purchased from Avanti Polar Lipids (Alabaster, AL, USA). Chloroform and methanol used for liposome preparation were purchased from Sigma-Aldrich Chemicals (Saint-Quentin-Fallavier, France). Fluorescent nucleotides used in this study were purchased from Jena Bioscience (https://www.jenabioscience.com/) under references NU-805-488 (for N^6^-(6-Aminohexyl)-ATP-ATTO-488) and NU-247-488 (for N^6^-(6-Aminohexyl)-ADP-ATTO-488). Fluorescent nucleotide analogs were diluted in 20 mM HEPES pH 7.5.

### Protein expression and purification

MreB^Gs^ was expressed in *Escherichia coli* and purified following a 2-step procedure with affinity purification (Ni-NTA agarose resin) and size exclusion chromatography. The protocol followed mainly the one previously described in Mao *et al*. (6). Two specific modifications were introduced for the production of protein for ssNMR: proteins eluted from the Ni-NTA affinity column dropped in 1 volume of buffer B (20 mM Tris pH 7, 500 mM KCl, 1 mM DTT, 2 mM EDTA), and size exclusion chromatography (HiLoad 16/60 Superdex 200 pg column from Cytiva) was pre-equilibrated and run with buffer C (Tris 20 mM; KCl 200 mM).

### Fluorescence anisotropy

Nucleotide exchange experiments were initiated by adding fluorescent nucleotides to MreB^Gs^ in buffer B (20 mM Tris pH 7, 500 mM KCl, 1 mM DTT, 2 mM EDTA) in presence or in absence of 5 mM CaCl_2_ or MgCl_2_. The binding of fluorescent nucleotide analogues to MreB^Gs^ was monitored by measuring changes in anisotropy using excitation and emission wavelengths of 504 nm and 521 nm, respectively. Steady-state values were recorded 5 min after the start of the experiment. The experiments were performed using a Safas Xenius XC spectrofluorometer (Safas Monaco) controlled by SP2000 software, version 7.8.13.0.

For binding affinity measurements, steady-state fluorescence anisotropy values of were measured at constant ATP* or ADP* concentrations in the presence of an increasing concentration of MreB^Gs^. Data were fitted and plotted using Igor Pro (version 9.0.5.1) to estimate K_d_ :

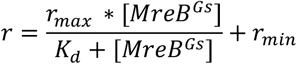

where r represents the fluorescence anisotropy readout, r_max_ represents the maximum anisotropy value when MreB^Gs^ is saturated with the fluorescent nucleotide, [MreB^Gs^] is the concentration of MreB^Gs^, K_d_ is the dissociation constant and r_min_ is the minimum anisotropy value corresponding to the fluorescent nucleotide alone in solution. Each data point represents the mean of at least three independent measurements, with error bars indicating the standard deviation. For kinetic experiments, the main reaction controlling the rate of the first nucleotide exchange (when MreB^Gs^ and ATP* or ADP* are mixed together) is the association of the fluorescent nucleotide with MreB^Gs^ (k_on,ATP*_ or k_on,ADP*_). The main reaction controlling the rate of the second nucleotide exchange (when an excess of ATP outcompetes ATP*) is the dissociation of the fluorescent nucleotide from MreB^Gs^ (k_off,ATP*_).

The two reactions involved are described by the following equations:

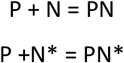

where P represents MreB^Gs^, N the unlabeled nucleotide, N_*_ the labelled nucleotide and PN and PN* the protein-nucleotide complexes. These equations involve four rate constants: k_on_^N*^, k_on_^N^, k_off_^N^ and k_off_^N*^.

The binding rate constants (k_on_) cannot be determined by this assay as they are too fast. However, the dissociation rates corresponding to the slowest reactions are accessible and can be mathematically described by the following differential equation:

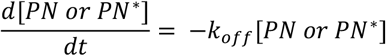

This equation can be resolved by fitting an exponential decay function to estimate the k_off_ value.

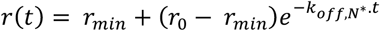

where r(t) represents the fluorescence anisotropy at time t, r_min_ is the minimum anisotropy value for the fluorescent nucleotide alone, r_0_ is the initial fluorescence anisotropy before adding the excess unlabeled nucleotide and k_off,N*_ is the dissociation rate constant of the fluorescent nucleotide.

### Liposome pelleting assays

Purified MreB^Gs^ was mixed at a final concentration (C_f_) of 1.34 μM (0.05 mg/mL) in a sedimentation buffer (Tris-HCl pH 7 20 mM, KCl 300 mM, MgCl_2_ 5 mM), in the presence or absence of ATP (C_f_ 2mM) and liposomes (C_f_ = 0.5 mg/mL). Liposomes of *E. coli* polar extract from Aventi (Merck) were prepared freshly by rehydrating desiccated lipids with H_2_O under strong vortexing. Mixtures were incubated for 1 h at room temperature prior to ultracentrifugation at 100 000*g* for 1 h at 18 °C. After careful recovery of the supernatants, the pellets were dispersed with an identical volume of sedimentation buffer supplemented with Tween20 at 0.1% final concentration. Aliquots of pellets and supernatants were loaded for analysis on a 12% SDS-PAGE, subsequently stained by Coomassie.

### Liposome preparation for AFM

Liposomes were prepared by dehydrating a lipid chloroform mixture of 20 mol % DOPG and 80 mol % DOPC in a desiccator for >3 hours at room temperature. Lipids were rehydrated with Citrate buffer (20 mM citrate pH 4.6 and 300 mM KCl). Liposomes were formed by vortexing and incubating at 40 °C. Lipid mixtures were passed 21 times through a lipid extruder with a 100 nm pore size on a heated plate at 50 °C (aimed internal temperature around 30°C) to produce small uniform-sized liposomes. A final concentration of 1 mM of liposomes was used. 3 mM of CaCl_2_ was added to the liposome suspension just prior to deposition.

### Nanofabrication of AFM cantilevers

BL-AC10DS-A2 cantilevers were purchased from Olympus (Tokyo, Japan) used as the scanning probe to visualize MreB polymerization. The spring constant (k) was 0.1 N/m while the resonance frequency (f) was 0.6 MHz in water (1.5 MHz in air). Its length, width, and thickness were 9, 2, and 0.13 μm, respectively. Electron-beam deposition (EBD) was performed to fabricate cantilever with a long and sharp tip, which is important to acquire high-resolution images. First, UV/O3 treatment was conducted to clean cantilevers, then the cantilevers were soaked in piranha solution (containing sulfuric acid and hydrogen peroxide). After that, EBD was carried out on the cantilevers (30 kV accelerating voltage, 2 min irradiation) using a field emission scanning electron microscope (ELS-7500, Elionix Inc., Tokyo, Japan).

### AFM nanoimaging

Atomic Force Microscopy nanoimaging was performed using a laboratory-built microscope, as previously reported (46–48). A laser beam with a 670 nm wavelength was focused on an EBD-processed cantilever tip through a 20x objective lens (CFI S Plan Fluor ELWD, Nikon, Tokyo, Japan). Dynamic cantilever deflection was detected by sensing the position of laser beam reflected by the cantilever with a position-sensing two-segmented photodiode. An optimal tapping force was generated by adjusting the free oscillation amplitude of the cantilever (A0) to 1.5–2.5 nm and the set point to 80–90% of the free amplitude. A glass stage glued with a stack of flat muscovite mica layer was attached to the HS-AFM scanner. Mica surface was treated with undiluted poly-L-Lysin for 10 min. The treated mica surface was washed with MilliQ and a liposome suspension of 20 mol % DOPG and 80 mol % DOPC (Avanti lipids) in Citrate buffer pH 4.6 with 300 mM KCl and 3 mM CaCl_2_ was added. After 25 min of incubation the Supported Lipid Bilayer (SLB) was washed with 25 mM HEPES pH 7.5. The experiment was performed in 25 mM HEPES pH 7.5, 100 mM KCL and 1 mM MgCl_2_. Purified His-MreB^Gs^ was centrifuged at 23 000 *g* for 10 min at 4°C to remove any precipitates. MreB was added to the SLB with or without 0.2 mM of ATP. Several locations of 0.7-1.3 µm^2^ were imaged on the SLB after 15 min to ascertain the presence and architecture of the filaments.

### MreB polymerization in the presence of membrane vesicles

Liposomes composed of ^2^H_31_-POPC, DOPC, and DOPG (35:45:20, w/w) were prepared by mixing lipid powders dissolved in chloroform/methanol (2:1 v/v). After solvent evaporation under a gentle air stream, the lipid film was rehydrated in 0.5 mL of Milli-Q water and lyophilized overnight. The resulting powder was resuspended in 100 µL of buffer A (20 mM Tris-HCl, pH 7.0, 200 mM NaCl, 1 mM DTT, and 2 mM EDTA) and homogenized by three cycles of vortexing, rapid freezing in liquid nitrogen (−196 °C, 1 min), and thawing at 40 °C for 10 min, yielding a milky MLV suspension. MreB in buffer A (23 μM) was added to preformed liposomes at a protein-to-lipid mass ratio 1:2, and incubated for 30 min at 30 °C. Next, when required, ATP and MgCl_2_ were added, and the mixture was incubated for another 10 min at 30 °C. The turbid proteoliposome suspensions (∼0.1-10 µm) were finally pelleted by ultracentrifugation (42,000 rpm, 1 h, 4 °C; MLA 80 rotor, Beckman Coulter).

### Solid-state NMR

^2^H NMR experiments were performed on a Bruker Avance III spectrometer operating at 77 MHz with a 4 mm double resonance HX-CPMAS probe. A phase-cycled quadrupolar echo sequence (90°x-t-90°y-t-acq) was used to perform the _2_H NMR experiments (49). Acquisition parameters were as follows: spectral window of 500 kHz, π/2 pulse width of 3.50 µs, interpulse delays (t) of 50 µs, recycle delays of 2 s for _2_H, and between 4096 and 20480 scans. Samples were equilibrated for 20 minutes at a given temperature before data acquisition. The temperature was regulated to ± 1 °C. Lorentzian line broadening was applied before Fourier transformation, starting from the top of the echo. To record undistorted^31^ P NMR spectra, we applied a Hahn spin echo sequence, 90°-t-180°-t-acq at the ^31^P frequency of 162 MHz on a 400 MHz (9.4 T) Bruker Avance III HD spectrometer (50), with a 90° pulse of 8 μs, an echo delay of 40 μs, a recycle delay of 5 s, a spectral window of 64 kHz, and a number of scans ranging from 1024 to 4096. All spectra were processed and analyzed using Bruker Topspin 4.0 software. Spectra were processed using a Lorentzian line broadening of 200 or 300 Hz for _31_P-NMR spectra.

### Computational analysis of ^2^H solid-state NMR spectra

Spectral processing was carried out in TopSpin 4.3.0 (Bruker), where an exponential window of 200 Hz line broadening was applied before performing the Fourier transform. De-Pake-ing (25, 51) and the calculation of the first-order spectral moment (M_1_) were carried out using NMR Depaker (provided by Dr. Sébastien Buchoux)(20); each M_1_ value was determined in triplicate, and the resulting standard deviations are shown as error bars in the plots. We assigned a default ±5% error to cover procedural uncertainties (26, 52).

To determine the S_CD_ order parameters along the ^2^H_31_-POPC acyl chain, we simulated wideline deuterium solid-state NMR spectra using a FORTRAN code developed by Erick Dufourc and accessed via a Microsoft.NET interface created by Arnaud Grélard. For each deuterated site along the deuterated lipid chain, we report the experimentally determined quadrupolar splitting (Δ*v*_Q_) and the count of deuterons at that specific acyl position based on the molecular structure of the lipid (Table S1). The simulations incorporated liposome deformation (reflected by a c/a parameter) to account for deviations from spherical bilayer normal distributions relative to the magnetic field (34). We adjusted the simulation to align with the experimental spectra through successive comparisons until optimal superposition was reached. From the best-fit simulation, we extracted the quadrupolar splitting for each labeled carbon and calculated the corresponding S_CD_ order parameters in the bilayer. We assigned a default ±5% error to all S_CD_ order parameters to cover both procedural uncertainties and intrinsic simulation variability; respecting the complexity of the system (three components), a safety margin chosen to exceed the experimental deviations reported in previous work (26) (52).

## Results

### MreB^Gs^ nucleotide-binding and dissociation kinetics

We first characterized the nucleotide-binding properties of MreB^Gs^ using fluorescent anisotropy, which is more sensitive and convenient for kinetic or rapid equilibrium studies than more traditional techniques such as isothermal titration calorimetry. A family of sensitive fluorescent nucleotides known as N^6^-(6-amino)hexyl-ATP derivatives was previously shown to bind to different types of eukaryotic actin, including muscle actin in mammals (11) and the distant cytoplasmic actin of the parasite *Leishmania* (12) while preserving nucleotide binding affinity and polymerization. We first determined whether these nucleotides also bind MreB^Gs^. The addition of N^6^-(6-amino)hexyl-ATP-ATTO-488 (hereafter referred to as ATP*) to MreB^Gs^ resulted in a rapid increase in fluorescence anisotropy (Fig. 1A). No change was observed in the absence of Mg^2+^ or Ca^2+^, indicating that pre-binding of divalent cations to MreB^Gs^ is essential for nucleotide binding (Fig. 1A), like in the case of actin (13). Purified MreB^Gs^ is presumably free of bound nucleotide in its active site since nucleotide binding was too fast to detect initial changes in anisotropy values with our instrument and thus could not be fitted accurately. However, we could measure plateau values that corresponded to steady-state binding (Fig. 1B). The higher plateau values observed in the presence of Mg^2+^ indicate that ATP* has a stronger affinity for Mg-bound MreB^Gs^ than for Ca-bound MreB^Gs^. Additionally, ATP* binding occurred more slowly in the presence of Ca^2+^, as steady state was reached at later time points than with Mg^2+^ (Fig. 1A). Based on these results, all subsequent experiments were conducted in the presence of Mg^2+^. We first quantified the binding affinity of ATP* by titrating increasing concentrations of MreB^Gs^ against a fixed concentration of ATP* (0.2 µM), and measuring fluorescence anisotropy at steady state. Binding curves revealed sub-micromolar dissociation constants (K_d,ATP*_ = 18 ± 5 nM; Fig. 1C). Identical measurements using the fluorescent equivalent ADP analogue (N^6^-(6-amino)hexyl-ADP-ATTO-488, ADP*) indicated that ADP binds MreB^Gs^ with similar -only slightly lower-affinity than ATP (K_d,ADP*_ = 27 ± 7 nM; Fig. S1A and Table 1). The K_d_ for both nucleotides is in the nanomolar range, like the reported K_d_ of ATP-monomeric actin (globular actin, G-actin) under physiological Mg^2+^ conditions (11, 13, 14) and indicating very high affinity. However, G-actin has a marked higher affinity for ATP than for ADP (15).

**Table 1.** Affinity and rate constants for nucleotide exchange on MreB^Gs^ measured in this study.

| Chemical reaction | $K_d$ (μM) | $k_{off}$ (s<sup>-1</sup>) |
| --- | --- | --- |
| Mg-MreB<sup>Gs</sup> + ATP* = Mg-MreB<sup>Gs</sup>-ATP* | 0.018 ± 0.005 | 0.0043 ± 0.0005 |
| Mg-MreB<sup>Gs</sup> + ADP* = Mg-MreB<sup>Gs</sup>-ADP* | 0.027 ± 0.007 | 0.0028± 0.0005 |

**Figure 1.**
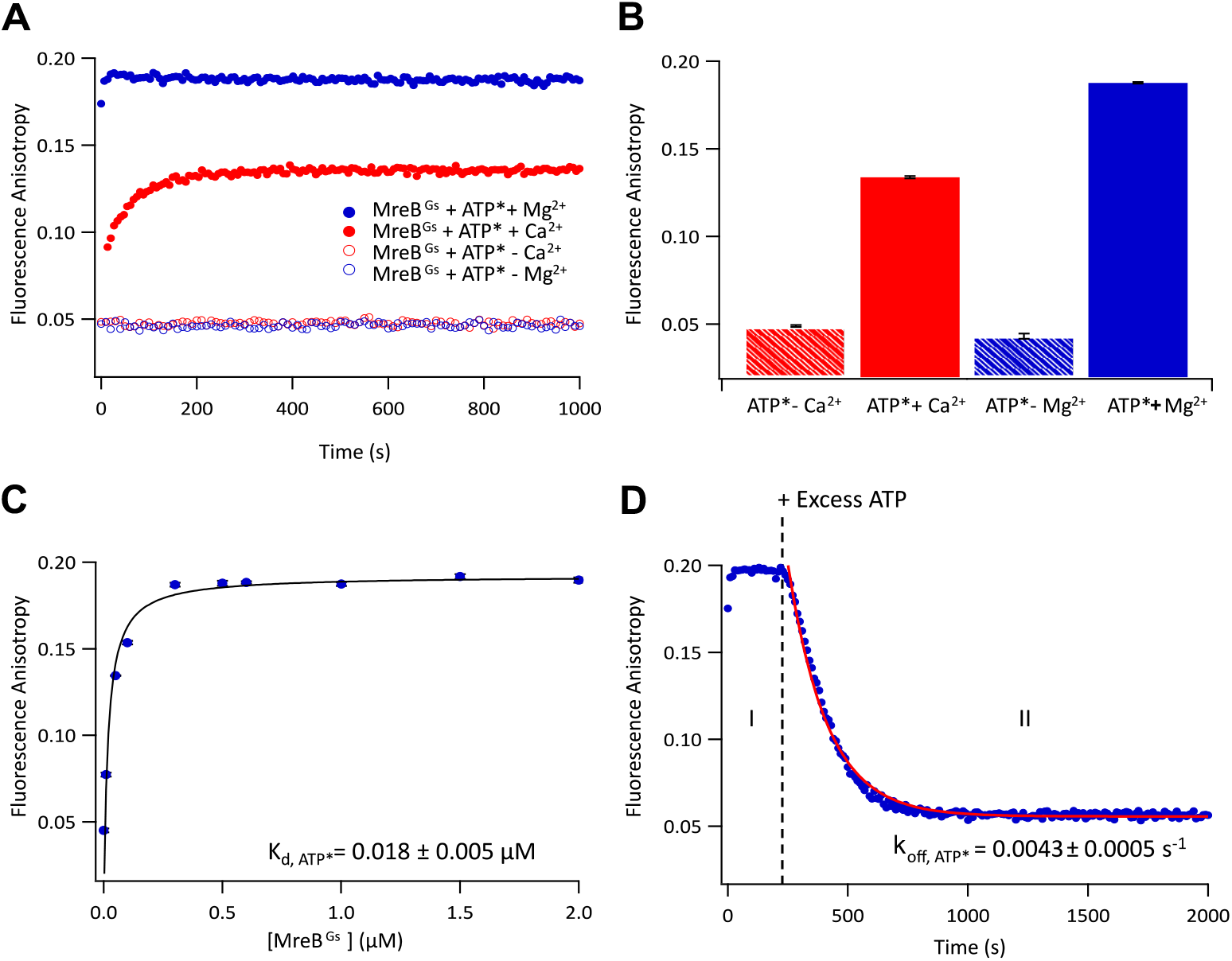
Analysis of MreB^Gs^–ATP binding and dissociation by fluorescence anisotropy. **A**. Binding kinetics of N6-(6-amino)hexyl-ATP-ATTO-488 (0.2 µM - referred to as ATP*) to MreB^Gs^ (1 µM), monitored by fluorescence anisotropy over time, in the presence (closed circles) or absence (open circles) of Ca^2+^ (red) or Mg^2+^ (blue) cations. **B**. Steady-state fluorescence anisotropy values of ATP* (0.2 µM) binding to MreB^Gs^ (1 µM) under the indicated ionic conditions. Bars represent mean ± standard deviation (n = 3 per condition). **C**. Fluorescence anisotropy measurements of ATP* binding to MreB^Gs^. Binding was assessed by titrating increasing concentrations of MreB^Gs^ against a fixed concentration of ATP* (0.2 µM), and measuring fluorescence anisotropy at steady state (5 minutes after mixing). Dissociation constants (K_d_) were determined by fitting the binding curves. Each data point represents the mean ± standard deviation from at least three independent measurements. **D**. Binding kinetics of ATP* (0.2 µM) to MreB^Gs^ (1 µM), followed in real time by fluorescence anisotropy. At 230 seconds, excess unlabeled ATP (1 mM) was added (dashed line), displacing bound ATP*. Red line: fitting of the dissociation phase (II) yields the k_off,ATP*_ value for ATP* release.

We next estimated the dissociation rates of both ATP* and ADP*. We pre-incubated MreB^Gs^ with the labelled nucleotide until equilibrium was reached, and triggered dissociation by addition of ∼200-fold excess unlabeled nucleotide (1 mM). The resulting decrease in fluorescence anisotropy was monitored over time and the dissociation curves of ATP* and ADP* from the MreB^Gs^-ATP* and MreB^Gs^-ADP* complexes, respectively, were fitted using a mono-exponential decay model (k_off,ATP*_ = 0.0043 ± 0.0005 s^−1^; k_off,ADP*_ = 0.0028 ± 0.0005 s^−1^ - Fig. 1D and S1B, respectively, and Table 1).

Taken together, these findings indicate that in the presence of Mg^2+^, MreB^Gs^ binds both ATP and ADP with very high affinity, indicating that under physiological conditions (*in vivo*) and at the experimental ATP concentrations used here, MreB^Gs^ is predominantly nucleotide-bound. Moreover, MreB^Gs^ exhibits only a slightly higher affinity for ATP* than for ADP*, yielding an ATP/ADP affinity ratio substantially lower than that observed for eukaryotic actin.

### ATP promotes polymerization of MreB^Gs^ on supported lipid bilayers

Though both ATP and ADP bind similarly to MreB^Gs^ (Table 1), only ATP was previously found to efficiently promote the formation of pairs of MreB^Gs^ filaments on lipid monolayers, as visualized by transmission electron microscopy (TEM), and the absorption of MreB^Gs^ onto lipid bilayers in QCMD experiments (6). We first confirmed the ATP requirement for binding to a lipid bilayer using a liposome pelleting assay. In the absence of liposomes, MreB^Gs^ remained mainly soluble, regardless of the presence of ATP (Fig. 2A). A fraction of MreB^Gs^ associated with liposomes in the absence of ATP, which may reflect the ion bridges-mediated intrinsic affinity of the protein for lipids. Consistent with this, increasing the concentration of salts (KCl), which reduces the contribution of electrostatic interactions to membrane binding by screening the charges of anionic lipid, abolished detectable binding of MreB^Gs^ to liposomes (Fig. S2). In the presence of both liposomes and ATP, most MreB^Gs^ sedimented as expected (Fig. 2A). We next turned to atomic force microscopy (AFM) to visualize MreB^Gs^ on a Supported Lipid Bilayer (SLB) in the presence and in the absence of ATP. We used SLBs made of dioleoylphosphatidylcholine (DOPC) doped with the anionic dioleoylphosphatidylglycerol (DOPG), which mimic *Bacillus* membranes and enable us to form SLBs on planar substrates (6). In the absence of added ATP, no MreB^Gs^ filaments were observed on the SLB (Fig. 2B). Some puncta could be seen, which could be either monomeric or low oligomeric MreB species, or small protein aggregates. In the presence of ATP, MreB^Gs^ rapidly formed a dense network of membrane-bound straight filaments (Fig. 2B), similar to what has been observed by TEM on lipid monolayers (6). Taken together, these findings confirmed that ATP promotes binding and polymerization of MreB^Gs^ on biomimetic lipid bilayers.

**Figure 2.**
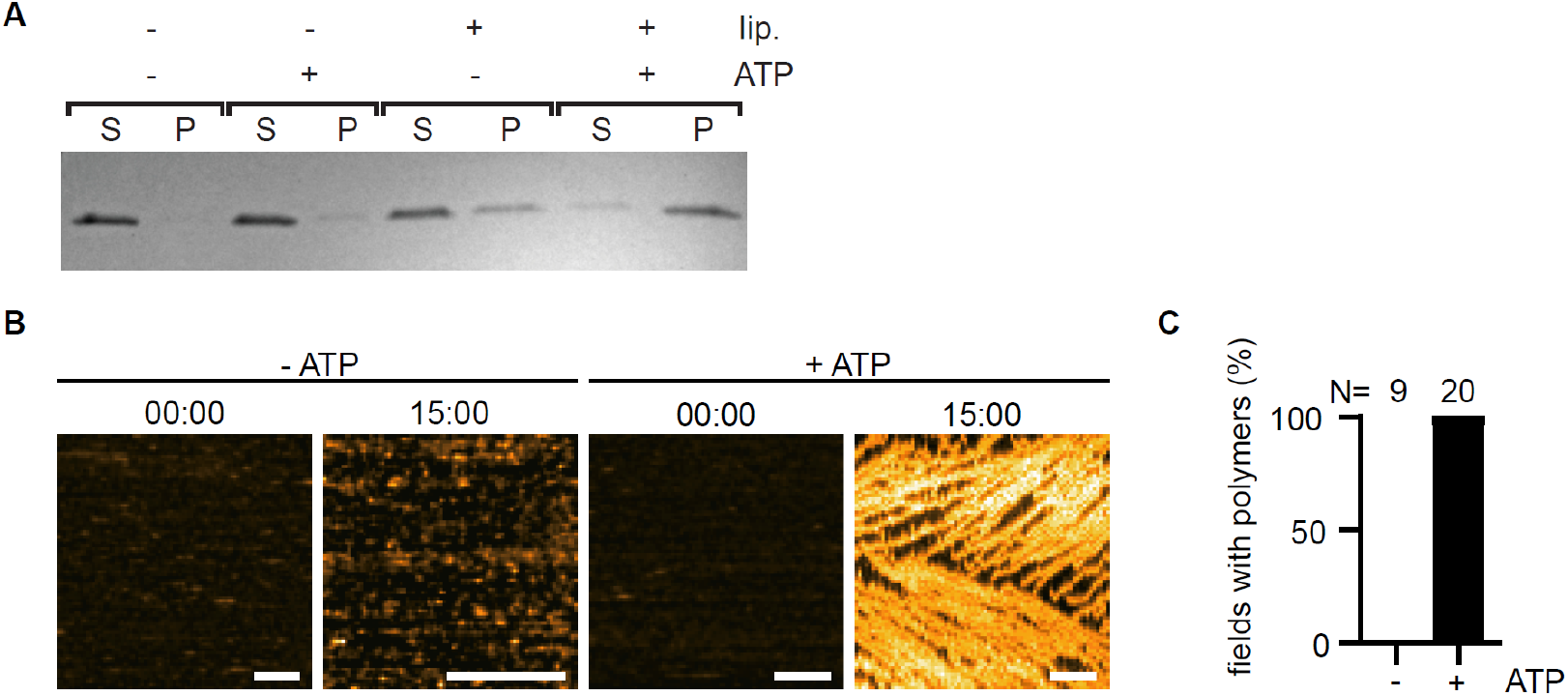
ATP-dependent binding and polymerization of MreB^Gs^ on lipid bilayers. **A**. Liposome pelleting assay. Representative 12% SDS-PAGE showing sedimentation of MreB^Gs^ (1.3 µM) in the absence and in the presence (2 mM) of ATP or liposomes (0.5 mg/ml). S, supernatant; P, pellet. **B**. AFM images of a SLB DOPC:DOPG 80:20 (molar ratio), previously shown to give a high absorption of MreB^Gs^ into SLBs (6), imaged before addition and 15 min after addition (min:sec) of MreB^Gs^ (1.2 µM) with or without 0.2 mM ATP. Scale bar, 200 nm. **C**. Quantification of the presence of filaments on AFM fields. N indicates the number of areas evaluated.

### ATP-dependent MreB^Gs^ interactions with the membrane detected by wideline ssNMR

We next monitored the interactions of ATP-bound MreB^Gs^ with the membrane using wideline ssNMR. Biological membranes can adopt different physical states, including solid-ordered gel, liquid-ordered, and liquid-disordered fluid phases, which differ in lipid mobility and packing. Wideline ssNMR is a precise, non-destructive method to probe lipid membrane mesophase, mostly a lamellar phase in the plasma membrane, membrane order, phase transitions and interactions under near-physiological conditions, typically detected on deuterium (^2^H) or phosphorus (^31^P) nuclei (16). Membrane properties are analyzed by measuring the residual quadrupolar splittings of deuterated lipid acyl chains (Δ*v*_Q_) or the chemical shift anisotropy of phosphorus headgroup (16–18). Because these anisotropic interactions are averaged out by motion, changes in lipid mobility translate into traceable variations in the observed ^2^H and ^31^P signals. Membrane multilamellar vesicles (MLVs) are pre-formed and tuned with respect to the lipid composition, allowing to reproduce a native membrane-mimicking environment. Subsequently, wideline ^2^H and ^31^P ssNMR report on local changes in mobility and interactions with molecular partners in the hydrophobic membrane core region and at the phosphorus headgroup, respectively. We doped vesicles of DOPC:DOPG 80:20 with _2_H-labeled POPC (palmitoyl-oleyl-phophatidyl-choline-d_31_) to produce ^2^H_31_-POPC:DOPC:DOPG (35:45:20 w:w) liposomes (Fig. 3A). To monitor the exclusive membrane interactions of polymerizing MreB^Gs^in comparison with soluble MreB^Gs^, we analyzed four analogously prepared samples of MLVs incubated with (1) buffer only, (2) buffer and MreB, (3) buffer and ATP, and (4) buffer with both MreB and ATP.

**Figure 3:**
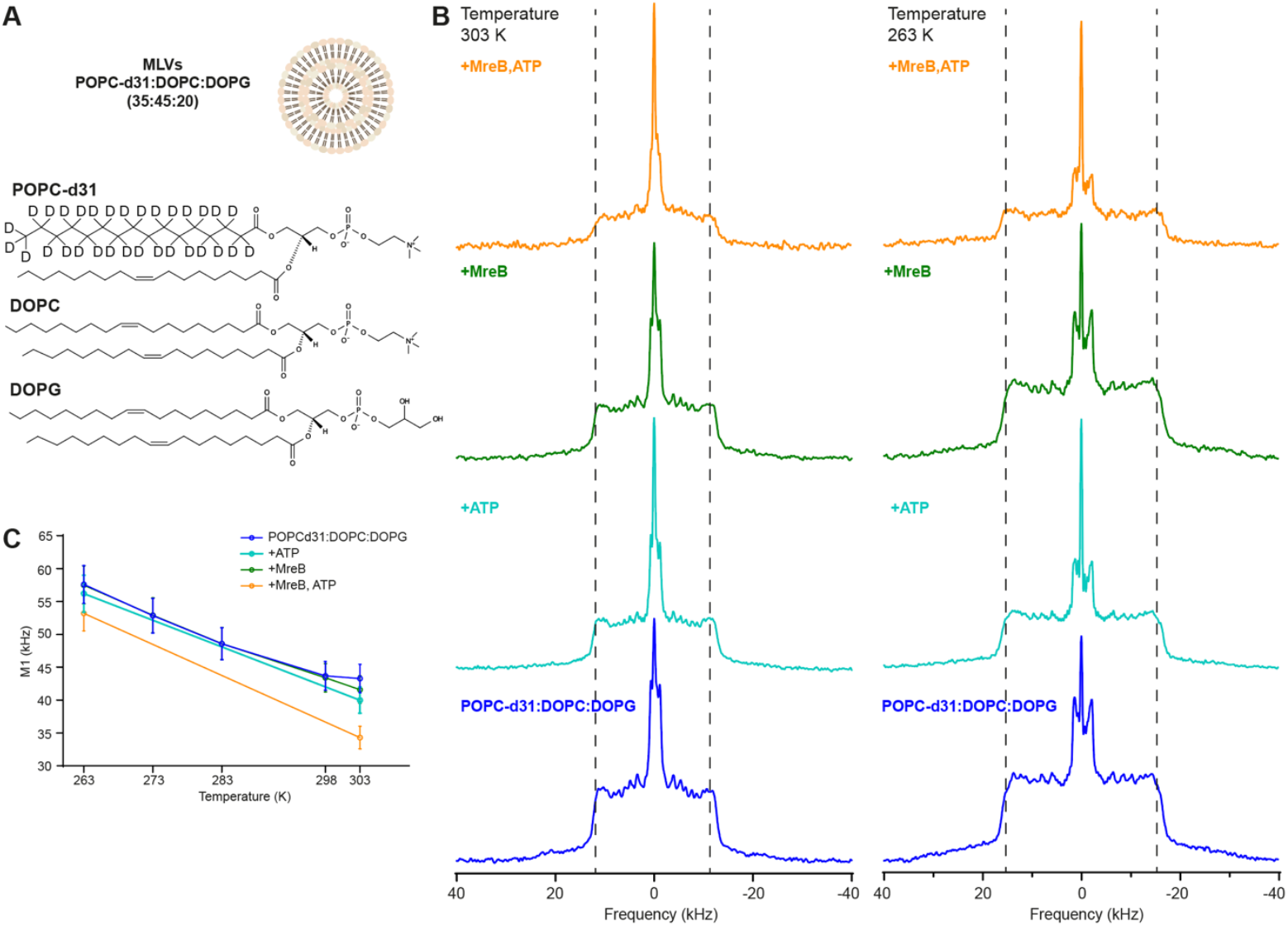
Effect of MreB on membrane dynamics. **A**. Chemical structures of ^2^H_31_-POPC, DOPC, and DOPG, used in a 35:45:20 mass ratio to reconstitute multilamellar vesicles. In ^2^H_31_-POPC, the palmitoyl chain is perdeuterated, with 14 methylene CD_2_ groups and one methyl CD_3_ group for ^2^H solid-state NMR. **B**. Comparison of ^2^H quadrupolar echo spectra of ^2^H_31_-POPC:DOPC:DOPG MLVs at 303 K and 263 K, recorded under buffer alone, ATP, MreB, and MreB+ATP conditions. Vertical dashed lines mark the regions corresponding to the largest quadrupolar splitting in the control condition. **C**. First spectral moment (M1) plotted as a function of temperature for multilamellar vesicles with buffer alone, ATP, MreB, or both MreB and ATP, at a protein-to-lipid mass ratio of 1:2. Error bars represent a ±5 % variation in M1 values due to methodological and measurement uncertainties.

We first recorded the quadrupolar splittings Δ*v*_Q_ (kHz) from the acyl chains of ^2^H_31_-POPC, which report on the order and mobility of different segments of the lipid tails (Fig. 3B). The largest splittings typically correspond to the plateau region of the first seven C-^2^H positions along the acyl chain, near the glycerol headgroup, which is sterically constrained and relatively rigid. The smaller splittings originate from the more flexible regions in the center of the membrane and from the terminal methyl group, which is the most disordered and gives rise to the narrowest doublet in the ssNMR spectrum. At 303 K, the ssNMR patterns indicate a fluid, liquid-disordered membrane, while at 263 K, closer to the fluid-to-gel phase transition temperature of the DOPC and DOPG acyl chains (Tm ≈ −16 °C), the lipid acyl chains become more ordered and rigid. In addition, a small central signal corresponds to ^2^HOH water molecules, detected at approximately 3 ppm (19). Together, these measurements allow us to characterize how lipid mobility varies along the acyl chain and how it changes with the temperature or in contact with a molecular partner.

The visual aspect of the signals recorded at 303 K were very similar for the conditions of phospholipid vesicles alone and in the presence of either ATP or MreB only, whereas the ^2^H spectral fingerprint was altered in presence of both MreB and ATP (highlighted by the dotted lines in Fig. 3B for eye guidance). We lowered the experimental temperature to 263 K, near the membrane phase transition, to examine the impact of MreB on the membrane in this altered membrane state. When comparing the spectra at 263 K, the three conditions of vesicles alone and with either ATP or MreB gave again similar spectra, in contrast to the spectrum of vesicles incubated with both MreB and ATP. To evaluate the impact of the interactions of polymerizing MreB with the membrane, we next calculated the first spectral moment M_1_ of the deuterated membrane lipids. This value represents an average of the order parameters along the lipid acyl chain and can be interpreted as variations in the mobility of the membrane phase (20–22). Higher M_1_ values correspond to more ordered, less mobile lipid chains, whereas lower values indicate increased chain flexibility and membrane fluidity. A significant reduction in the global order of the membrane hydrophobic core was observed in the presence of both MreB and ATP, suggesting clear interactions between MreB filaments and the membrane as well as a local fluidifying effect (Fig. 3C). The membrane disordering effect was maintained when shifting the membrane to lower temperatures approaching the membrane fluid-to-gel phase transition (Fig. 3C), potentially due to a continuous local membrane destabilizing effect of MreB filaments, favoring the fluid liquid-disordered membrane phase.

### MreB^Gs^ filaments promote fluidization of the lipid bilayer core

We next calculated the carbon-deuterium order parameters (S_CD_) at each carbon position of the lipid acyl chains using the experimentally measured Δ*v*_Q_. The S_CD_ value describes how much the residual quadrupolar couplings are averaged by the mobility of the C-D bond at each position of the acyl chain with respect to the bilayer normal (16, 22). In other words, it provides a direct measure of how restricted in motion each segment of the lipid chain is on the sub-microsecond timescale. To obtain the S_CD_, the quadrupolar splitting measured at each carbon position k (Δ*v*Q^k^) is converted into an order parameter using the relation Δ*v*Q^k^ = 0,75*A_Q_ S_CD_^k^, where A_Q_ is the static quadrupolar coupling constant (167 kHz) (22–24). Order parameter of the acyl chain typically vary between disordered (2*|S_CD_| = 0) to fully rigid systems (2*|S_CD_|= 1). To interpret our experimental data, we simulated the expected _2_H NMR spectra using established algorithms. These simulations take into account both the ‘de-paked’ oriented-like spectra (20, 21, 25) as well as the MLV ellipsoidal deformation in the magnetic field represented by the c/a value (where c represents the axis of the ellipsoid parallel to the magnetic field). The spectra showed no significant ellipsoidal deformation of the vesicles in the magnetic field under any of the four tested conditions (Fig. 4A). We also observed the common S_CD_ plateau region (22, 26), ranging from carbon position 2-7, for which S_CD_ values remain rather constant (Fig. 4B, Table S1), reflecting the sterically constrained, more rigid carbon-deuterium bonds near the phospholipid glycerol backbone. Neither ATP nor MreB alone affected the order parameter S_CD_ of the acyl chain relative to the buffer-only control. However, MreB in the presence of ATP induced a clear reduction of the S_CD_ considering the carbon positions 8 to 16, corresponding to the part of the acyl chains located in the inner, hydrophobic region of the lipid bilayer (Fig. 4B, Table S1). This destabilizing effect, reflected by increased mobility of the membrane core acyl chains, likely arises from peripheral protein segments inserting into the lipid headgroup region, perturbing lipid packing and generating additional space for acyl chain fluctuations. Similar effects have been observed for membrane-binding peptides (27, 28).

**Figure 4.**
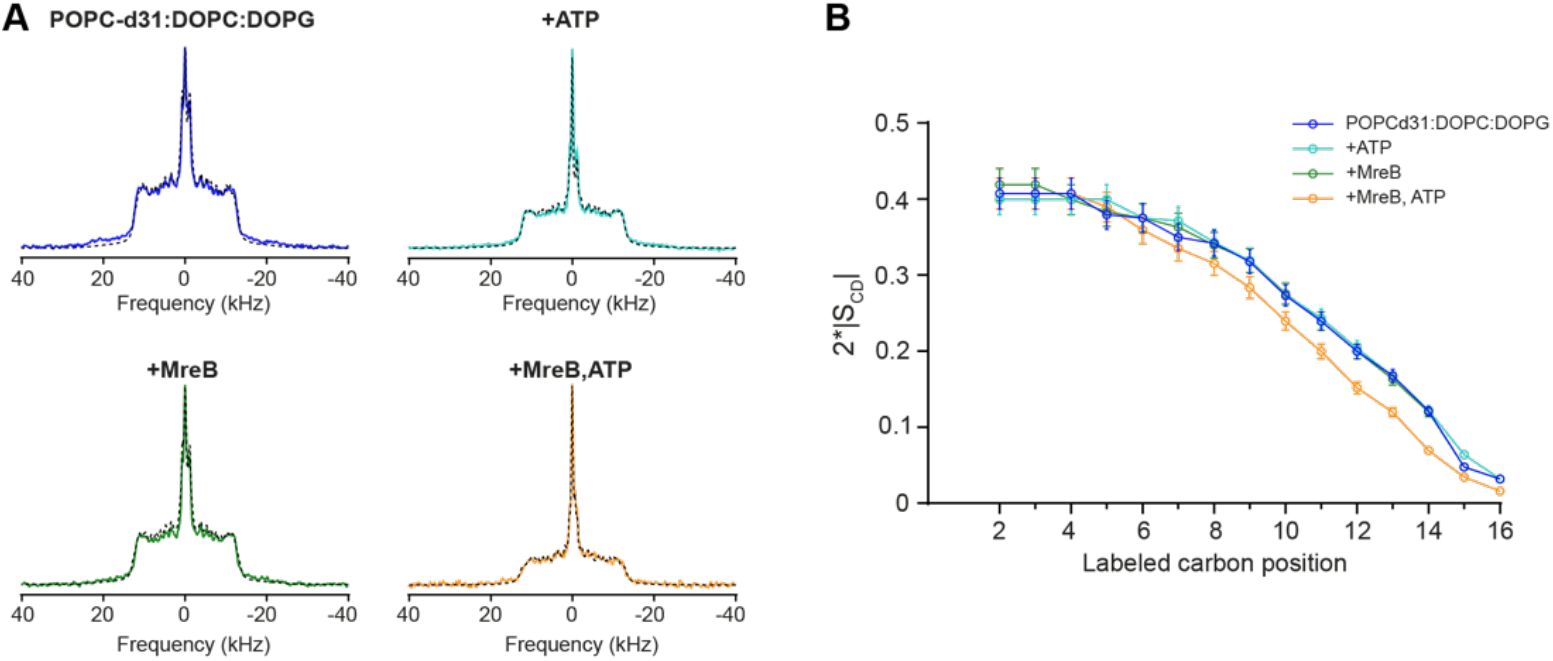
Impact of MreB in membrane order. **A**. Spectral simulations (black dashed lines) overlaid with the experimental ^2^H quadrupolar echo spectra acquired at 303 K under the four experimental conditions: buffer alone, ATP, MreB, and MreB + ATP. **B**. Carbon-deuterium order parameters 2*|S_CD_| of ^2^H_31_-POPC in ^2^H_31_-POPC:DOPC:DOPG multilamellar vesicles (35:45:20) at 303 K, measured in the four experimental conditions. Error bars indicate a ±5% uncertainty in S_CD_ values, reflecting both methodological and simulation-related variability.

### ssNMR shows association of ATP-MreB to the lamellar membrane phase, with no detectable Pi release

We next sought to characterize the nucleotide-bound state of ATP-induced MreB^Gs^ filaments on the preformed lamellar membrane. Previous results using non-hydrolysable analogs of ATP suggested that ATP hydrolysis was required for MreB^Gs^ membrane binding and polymerization (6). In agreement with this, Pi release by ATP-bound MreB^Gs^ was favored in the presence of lipids (6). We turned to ^31^P-detected wideline ssNMR under static conditions and analyzed the nucleotide phosphorus shifts and the residual chemical shift anisotropy (CSA) of phosphorus in the phospholipid headgroups. The CSA reflects the orientations and dynamics of the phosphate groups, providing information about the bilayer phase and its structural organization.

Across the four experimental conditions tested, the ^31^P NMR line shapes were consistent with a lamellar phase (17, 26) (Fig. 5). In a lamellar membrane phase, lipid molecules arrange in flat, parallel sheets (lamellae) that are typical biological membrane arrangements (29) and at the origin of a characteristic phosphorus spectral envelope (30). The CSA was measured from the width of the ^31^P signals and reflects the range of orientations of the phosphate groups relative to the magnetic field. From our spectra, we obtained a CSA of approximately 49 ppm (8 kHz), in agreement with previous reports (17, 26) (Fig. 5). Addition of MreB^Gs^ alone introduced minor contributions of an isotropic peak around 0 ppm (Fig. 5), reflecting the orientation averaging of the phospholipid headgroup, as in occurs in rapid tumbling lipids or micelles, potentially generated through electrostatic interactions of MreB with the lipids. Upon the addition of either ATP alone or MreB + ATP, the interpretation was complexified due to the superposition of ^31^P signals originating from free ATP and bound ATP (or a post-hydrolytic ADP-P_i_ species). Notably, the ^31^P signals of ATP corresponding to the α, β and γ positions were all detected but shifted by 1 ppm, 0.9 ppm and 0.8 ppm, respectively, in the presence of MreB (Fig. 5 and S3, Table S2), and the CSA pattern still reflected a lamellar membrane phase. We cannot exclude that only the resonances of the α and β positions shifted while, in this case, the γ resonance remained nearly unchanged. No isotropic signal expected for free, rapidly tumbling ^31^P (typically observed around 3.7 ppm in water (31)) was observed in the MreB + ATP sample (Fig. 5), suggesting that no continuous ATP hydrolysis occurred and thus no detectable Pi was released from ATP-bound filaments under these conditions.

**Figure 5.**
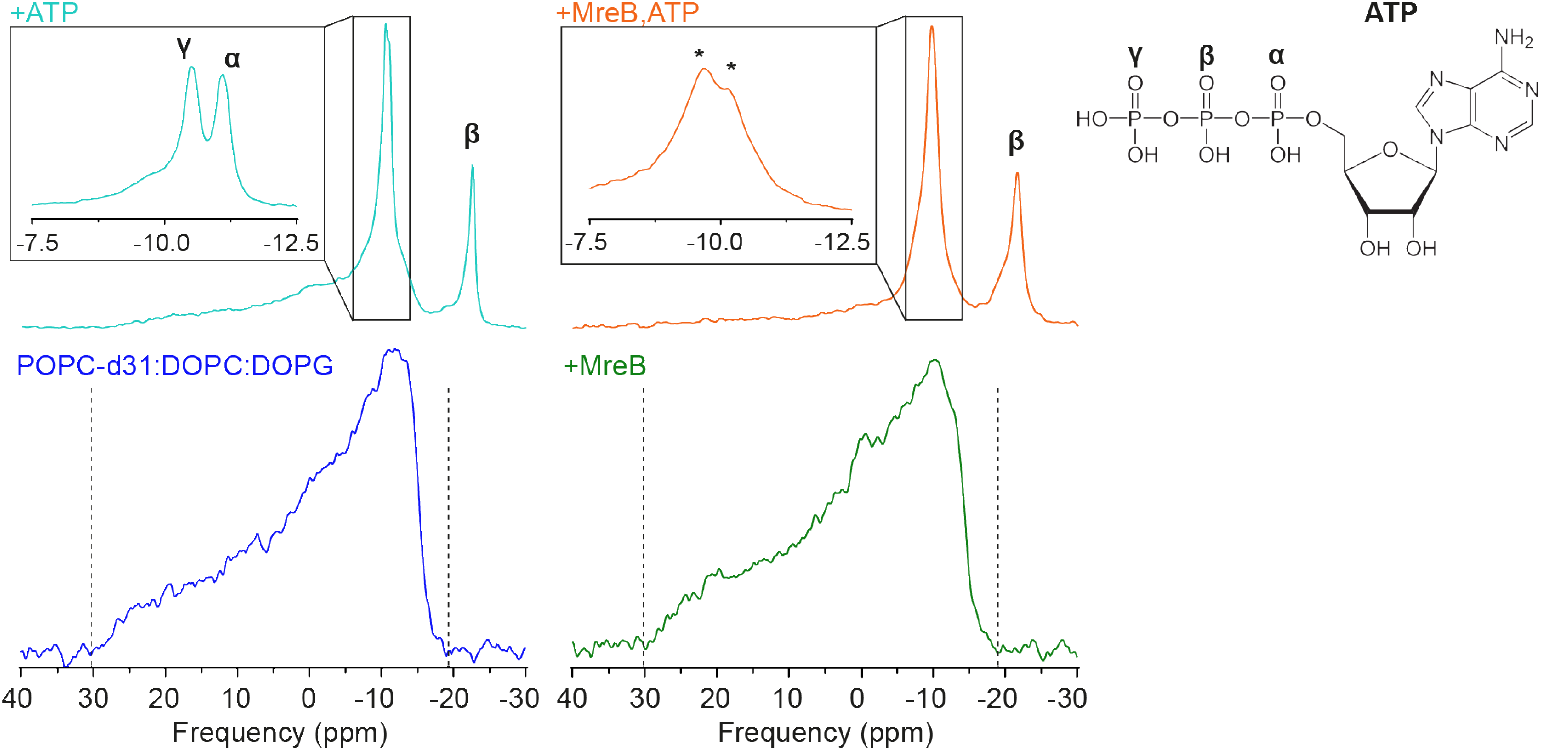
Evaluation of MreB^Gs^-induced changes in phosphorus properties of lipid headgroups and ATP. ^31^. P NMR spectra of ^2^H_31_-POPC:DOPC:DOPG multilamellar vesicles (35:45:20) recorded at 303 K in the presence of buffer alone (blue), supplemented with ATP alone (cyan), MreB^Gs^ alone (green), or with both MreB^Gs^ and ATP (orange). Signals resulting from the three phosphate groups of ATP are indicated by the corresponding Greek letters. Magnified regions (boxed) are shown to discriminate the γ and α phosphate signals of ATP. In spectra recorded in the presence of both MreB^Gs^ and ATP (orange), the α-phosphate and the γ-phosphate positions are indicated with an asterisk, as the resonances may derive from the α and γ in ATP or in trapped ADP+Pi due to ATP hydrolysis (6).

## Discussion and conclusion

In this study, we used three highly sensitive techniques: fluorescence anisotropy, AFM and ssNMR, to characterize the binding of ATP to MreB^Gs^ and the effect of MreB binding to a membrane surface with unprecedented resolution. Our results reveal a high-affinity interaction between ATP and MreB^Gs^, which is essential for filament formation on lipid bilayers, resulting in disruption of lipid packing and local fluidization of the membrane. While ssNMR analysis of ^2^H quadrupolar splitting is standardly used to probe membrane binding of peptides or proteins as well as changes in membrane phase and order (26–28, 32–34), to our knowledge this is the first report demonstrating its ability to detect nucleotide-induced membrane-binding and lipid-disordering by a polymerizing protein, highlighting its exceptional sensitivity to detect changes in the chemical environment and its applicability.

In the absence of ATP, a small fraction of MreB^Gs^ co-pellets with liposomes in our liposome-binding assays. Increasing the salt concentration abolishes this association, likely by screening membrane negative charges, indicating that the interaction is electrostatic. Consistent with this, a very small fraction of lipids appears to be extracted from the lipid bilayer, potentially through electrostatic contacts between monomeric MreB^Gs^ and the membrane surface, resulting in an isotropic signal close to the noise level in the ^31^P spectra. However, neither the membrane order nor the lamellar phase or phosphorus headgroup environment is perturbed, indicating the absence of significant protein-membrane interactions. This is further supported by the close similarity of the ^2^H-detected ssNMR spectra obtained with vesicles and with vesicles incubated with MreB^Gs^ alone. Taken together, these data suggest that monomeric MreB^Gs^ has peripheral contact with the membrane through electrostatic interactions, but does not insert into the hydrophobic core of the bilayer.

In the presence of ATP, the high affinity and rapid binding kinetics of MreB^Gs^ for the nucleotide ensure that, as observed for actin, MreB^Gs^ rapidly becomes fully ATP-bound and ready for polymerization. ATP-bound MreB^Gs^ assembled into filaments on SLBs as shown by AFM, and induced clear changes in the ^2^H-detected ssNMR spectra, revealing membrane interactions that likely result from the insertion of peripheral protein segments into the lipid bilayer. Peptides and proteins can exhibit a varying effect on membrane order and thereby lipid chain mobility (26–28, 32, 34, 35). Our data indicates that the disordering effect of ATP-bound MreB^Gs^ on acyl chain dynamics is moderate but significant.

The membrane insertion region likely corresponds to the hydrophobic spatially close N-terminus and α2-β7 loop of MreB^Gs^, which have been shown to be required for lipid interaction (6). We previously proposed that the transition from soluble MreB^Gs^ to membrane-bound polymers could be driven by a structural change triggered by ATP hydrolysis, which would render this region prone for insertion in the membrane (6). We did not detect an isotropic free phosphate signal in our ^31^P ssNMR spectra of MreB^Gs^ in the presence of ATP, which would be expected if ATP hydrolysis was required for membrane binding and free Pi was continuously released from the membrane-bound filaments, as previously reported (6). Although it cannot be formally excluded, it is unlikely that a large part of MreB-bound ATP was hydrolyzed to ADP and all Pi molecules were lost in the ssNMR ultracentrifugation step. The lack of detectable free Pi in our ssNMR experiments, while AFM showed a dense network of membrane-bound filaments in similar biochemical conditions, rather suggests absence of significant ATP hydrolysis, or else trapping of the hydrolyzed Pi in the nucleotide-binding cleft during the course of the ssNMR experiment. Binding of either ATP or ADP-Pi-to the nucleotide-binding pocket would be consistent with the ^31^P chemical shift changes observed for all three ATP phosphate groups relative to unbound ATP, indicating a different chemical environment. The ADP-Pi-bound intermediate was previously suggested to be the long-lived nucleotide-bound form within MreB filaments (6). It cannot be ruled out, however, that ATP was the form detected in our ^31^P ssNMR spectra, and thus that ATP hydrolysis is not strictly required for membrane binding and polymerization, but occurs later, within membrane-bound MreB filaments, as is the case for actin filaments. Under this alternative model, ATP binding itself, rather than ATP hydrolysis, would trigger the structural change involved in the switch from the soluble to the membrane-affine form. Interestingly, while the crystal structure of the ATP-bound form and the nucleotide-free (apo) form of MreB^GS^ (PDB ID 8AZG and 7ZPT, respectively) are highly similar (6), ATP binding induces changes in the two membrane anchoring domains (Fig. S4). The end of the N-terminal region is predicted to be intrinsically disordered and was not resolved in the crystal structures (aa 1-5 and 1-3 missing, respectively). However, both the α2-β7 loop and the segments of the N-terminal extremity present in the crystals have a high B-factor (also known as the atomic displacement parameter) in the ATP-bound form of MreB^Gs^ (Fig. S4), which reflects a higher disorder and likely a higher flexibility (36), favoring structural adaptations such as membrane insertion.

While MreB^Gs^ does not polymerize on a lipid surface in the presence of ADP (6), our anisotropy data show that soluble MreB^Gs^ binds both ATP and ADP with comparable affinities under physiological Mg^2+^ conditions, with affinity for ATP being less than twofold higher than for ADP. In contrast, G-actin has a markedly higher affinity for ATP versus ADP, about 4-fold under physiological Mg^2+^ conditions (and 200-fold in the presence of Ca^2+^) (14, 15). Both eukaryotic and bacterial cells have similar (high) millimolar intracellular ATP concentrations (average in the 1-5 mM range) and maintain a similar (high) ATP/ADP average ratio of ∼ 5-10 (37–39). Because of actin’s substantially greater affinity for ATP compared to ADP, coupled with the prevalence of G-actin bound to profilin, an essential actin binding protein (ABP) that catalyzes ADP-to-ATP exchange (13), the high majority of monomeric actin in eukaryotic cells is predicted to exist in the ATP-bound, polymerization-competent state. In bacterial cells, which lack known counterparts of ABPs, the similar affinities between MreB^Gs^ and both ATP and ADP found here suggest that the fraction of ATP-bound MreB^Gs^ is substantially lower, with a non-negligible fraction of monomeric MreB^Gs^ bound to ADP. Although the intracellular ATP/ADP ratio ([ATP] = 2-2.6 mM and [ADP] = 0.4-0.8 mM measured in *Escherichia coli* (39, 40)) would imply that most MreB^Gs^ monomers are ATP-bound, this ratio is known to vary depending on growth conditions such as nutrient availability, growth rate, or environmental stress. Thus, the similar affinities of MreB^Gs^ for ATP and ADP may provide a mechanism for modulating the pool of polymerization-competent MreB in bacteria.

Finally, our data show that MreB^Gs^ in polymerizing conditions impacts on the membrane stability, locally increasing the mobility of the acyl chain in the membrane inner core. In line with this, recent molecular dynamics simulations suggest that *E. coli* MreB filaments induce local membrane bending by lipid displacement and physical distortion via the N-terminal amphipathic helix of *E. coli* MreB, which mediates membrane binding (41). Both shallow insertion of membrane-anchoring segments and membrane bending are mechanisms that can increase lipid chain mobility and thus fluidify membranes. Our finding that MreB^Gs^ filaments fluidify the membrane is in agreement with previously reported *in vivo* data that suggested that MreB filaments generate fluid lipid domains (regions with increased fluidity, RIFs) in the *B. subtilis* plasma membrane (10). Thus, in Gram-positive bacteria at least, membrane-associated MreB filaments may directly contribute to local fluidization of the phospholipid membrane. Interestingly, AFM imaging demonstrated that the membrane-bound cell wall building block Lipid II co-separates in the liquid-crystalline (also known as liquid-disordered) more fluid regions of SLBs (42). In agreement with this, Lipid II as well as the Lipid II synthase MurG were reported to preferentially colocalize with RIFs in *B. subtilis* (43, 44). Although the mechanisms behind the fluidifying effect measured *in vitro* might differ from those at the origin of the effect observed *in vivo*, it is tempting to speculate that increased membrane fluidity by membrane-bound MreB filaments might locally increase the concentration of Lipid II and/or modulate the activity of integral membrane proteins of the Rod complex (45), by increasing the rates of conformational changes or protein-protein or protein-substrate interactions. Future studies will be required to explore whether local, MreB-dependent, membrane fluidization influences the activity of the cell wall elongation machinery.

## Data availability

Research data will be deposited on Zenodo after publication or be available from the corresponding authors upon request.

## Authors contributions

I.E.A., A.A.M, C.D., E.M, K.S.L., A.C. performed the research. I.E.A., A.A.M., C.D., E.M., E.J.D., B.H. and A.C. analysed data. R.W., A.M, R.C.-L. and B.H. provided infrastructure and scientific supervision. R.C.-L. and B.H. designed the research, funded the research and wrote and revised the manuscript, with input from all authors.

## Acknowledgments

We thank Lukas Kalvoda and Martin Lenz, along with members of the ProCeD lab, for helpful discussions. This project has received funding from the European Research Council (ERC) under the Horizon2020 research and innovation program (ERC CoG No 772178 to R.C.-L.), the French National Research Agency (ANR-23-CE11-0005-01 to BH), a Kanazawa University WPI-NanoLSI Bio-SPM collaborative research grant, and the Bordeaux University Health sciences and technologies department. This study has also received financial support from the French government in the framework of the France 2030 programme IdEx université de Bordeaux, and benefited from the facilities and expertise of the Biophysical and Structural Chemistry platform at IECB, CNRS UAR3033, Inserm US001, Univ. Bordeaux.

