## Supplementary material for "ATP-driven membrane binding and polymerization of bacterial actin MreB promotes local membrane fluidization": Supp. material

### Supplemental information

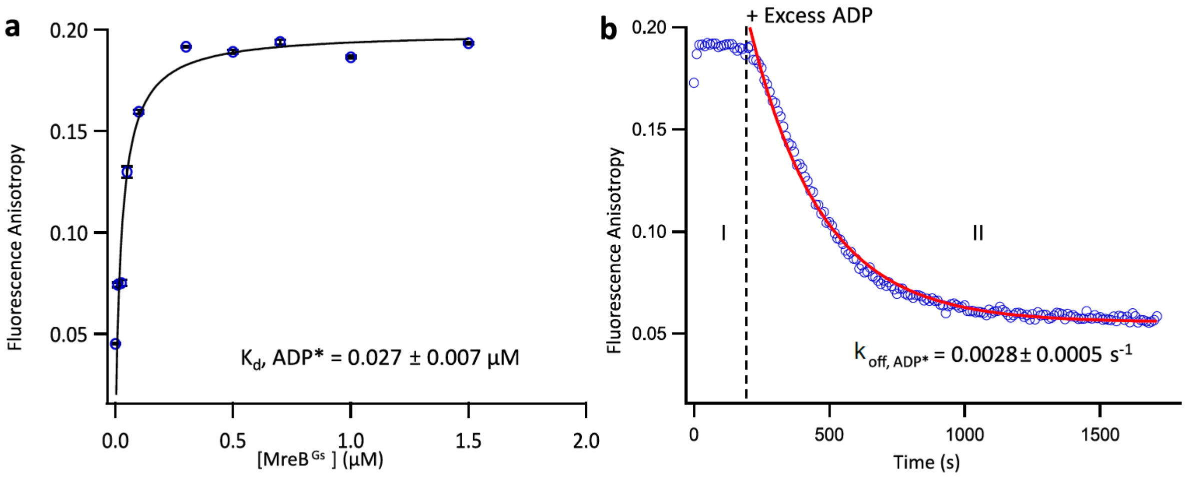


**Supplementary Figure 1. Analysis of MreB$^{Gs}$–ADP* binding by fluorescence anisotropy. A**. Fluorescence anisotropy measurements of N6-(6-amino)hexyl-ADP-ATTO-488 (0.2 µM - referred to as ADP*) to MreB$^{Gs}$ (1 µM). Binding was assessed by titrating increasing concentrations of MreB$^{Gs}$ against a fixed concentration of ADP* (0.2 µM), and measuring fluorescence anisotropy at steady state (5 minutes after mixing). Dissociation constants ($K_d$) were determined by fitting the binding curves. Each data point represents the mean ± standard deviation from at least three independent measurements. **B.** Binding kinetics of ADP* (0.2 µM) to MreB$^{Gs}$ (1 µM), followed in real time by fluorescence anisotropy. At 230 seconds, excess unlabeled ADP (1 mM) was added (dashed line), displacing bound ADP*. Red line: fitting of the dissociation phase (II) yields the $k_{off,ATP*}$ value for ADP* release.

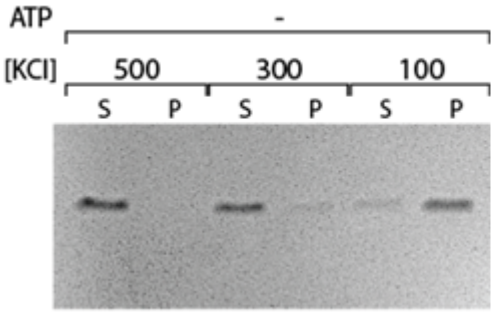


**Supplementary Figure 2. The Ionic strength affects MreB$^{Gs}$ binding to lipids in a liposome pelleting assay.** 12% SDS-PAGE showing sedimentation of MreB$^{Gs}$ (1.3 µM) in the presence of *E. coli* lipid polar extract liposomes (0.5 mg/ml) in the absence (-) of ATP. The reactions contain either 100, 300 or 500 mM KCl. S, supernatant; P, pellet.

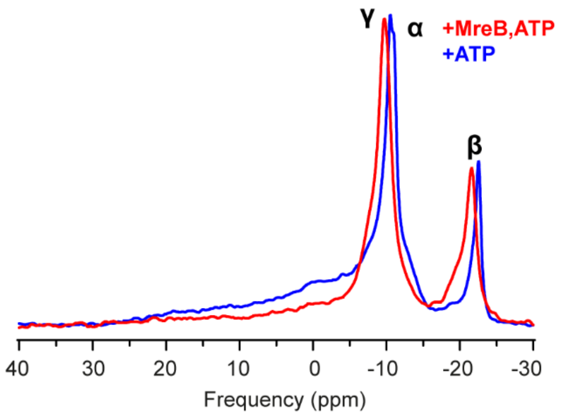


**Supplementary Figure 3.** Superimposition of $^{31}P$ NMR spectra of $^{2}H_{31}$-POPC:DOPC:DOPG multilamellar vesicles (35:45:20) recorded at 303 K in the presence of ATP (blue) and MreB + ATP (red), showing the assignments of the three phosphate groups of ATP.

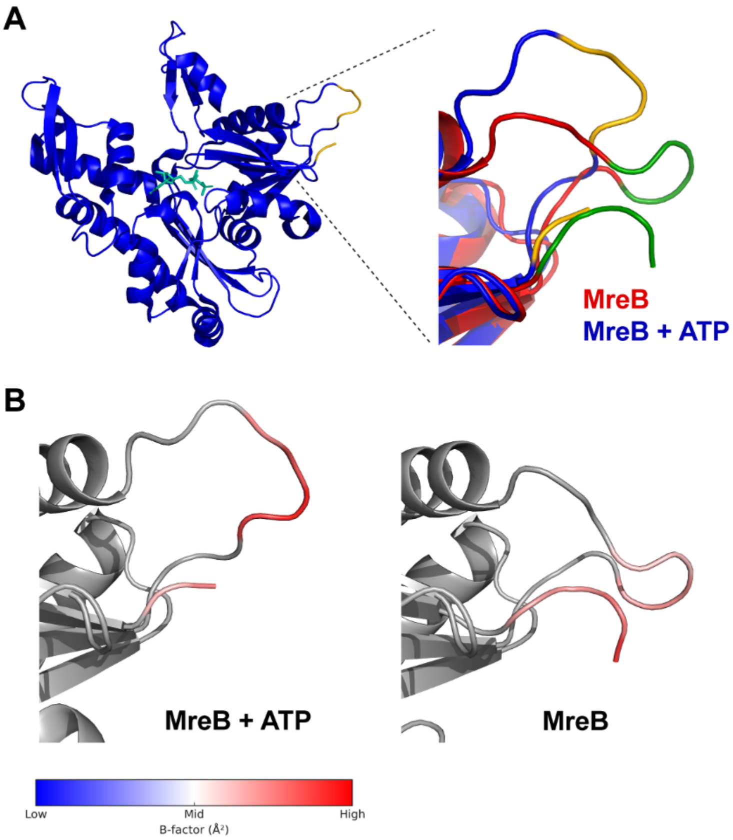


**Supplementary Figure 4. Local flexibility in the membrane-binding region of apo and ATP-bound MreB. A.** Left: Crystal structure of ATP-bound MreB from *G. stearothermophilus* (PDB ID 8AZG), highlighting the partial N-terminal region and the α2-β7 loop (yellow), which are essential for membrane binding and polymerization on lipid surfaces. Right: zoom in and structural overlay of apo MreB$^{Gs}$ (PDB ID: 7ZPT, red, with the N-terminal region and α2-β7 loop highlighted in green) and ATP-bound MreB$^{Gs}$ (PDB ID: 8AZG, blue) within the membrane-binding region, showing a conformational shift associated with nucleotide binding. **B.** Zoom up representation of apo MreB (PDB ID: 7ZPT) and ATP-bound MreB (PDB ID: 8AZG), showing the N-terminal segment and the α2-β7 loop, which are highlighted and colored according to their B-factor values, which were extracted from the PDB files using PyMOL and averaged per residue. Higher B-factors reflect greater atomic mobility and flexibility, highlighting regions with increased dynamic behavior upon nucleotide binding.

**Supplementary Table 1**: Simulated quadrupolar splittings $\Delta\nu_Q$ of the $^2H_{31}$-POPC acyl chain in $^2H_{31}$-POPC:DOPC: DOPG (35:45:20, w/w) vesicles under the four experimental conditions.

| **Deuterated carbon position** | **Experimental condition** | | | |
|---|---|---|---|---|
| | **Buffer** | **Buffer + ATP** | **Buffer + MreB** | **Buffer + MreB + ATP** |
| | $\Delta\nu_Q$ | $\Delta\nu_Q$ | $\Delta\nu_Q$ | $\Delta\nu_Q$ |
| **2** | 51 | 50 | 52.5 | 51 |
| **3** | 51 | 50 | 52.5 | 51 |
| **4** | 51 | 50 | 50 | 51 |
| **5** | 47.5 | 50 | 48 | 48.8 |
| **6** | 47 | 47 | 47 | 45 |
| **7** | 43.8 | 46.5 | 45.5 | 42 |
| **8** | 42.8 | 43 | 42.5 | 39.5 |
| **9** | 39.8 | 40 | 40 | 35.5 |
| **10** | 34.2 | 34.3 | 34.5 | 30 |
| **11** | 30 | 30.5 | 30 | 25 |
| **12** | 25 | 25.5 | 25 | 19 |
| **13** | 21 | 21 | 20.5 | 15 |
| **14** | 15.2 | 15.3 | 15 | 8.7 |
| **15** | 6 | 8 | 6 | 4.3 |
| **16** | 4 | 4 | 4 | 2 |

**Supplementary Table 2**: ATP $^{31}P$ chemical shifts at positions α, β and γ.

| **Sample** | **$^{31}P$ chemical shift ATP (ppm)** | | |
|---|---|---|---|
| | **α** | **β** | **γ** |
| **ATP** | -11.1 | -22.6 | -10.5 |
| **MreB + ATP** | -10.1 | -21.7 | -9.7 |
